# Multi-parametric NIR-II fluorescence perfusion imaging towards quantitative stroke evaluation

**DOI:** 10.64898/2026.09.02.749025

**Authors:** Liangtao Gu, Zhao Fang, Chen Duan, Yu Zhang, Xieyang Bian, Xinning Sun, Jiale Wang, Peixi Peng, Yanan Wu, Tinghua Wen, Guoyan Zheng, Hao Chen, Wuwei Ren

**Author notes:** Correspondence to: Wuwei Ren, School of Information Science and Technology, ShanghaiTech University, Shanghai 201210, China; Hao Chen, State Key Laboratory of Chemical Biology, Molecular Imaging Center, Shanghai Institute of Materia Medica, Chinese Academy of Sciences, Shanghai 201203, China; Guoyan Zheng, Institute of Medical Robotics, School of Biomedical Engineering, Shanghai Jiao Tong University, Shanghai 200030, China. Equal Contributions (Liangtao Gu, Zhao Fang, Chen Duan). Author contributions: Wuwei Ren, Hao Chen and Guoyan Zheng conceived the study. Liangtao Gu, Chen Duan, Jiale Wang, Peixi Peng, and Tinghua Wen developed the algorithm. Chen Duan and Zhao Fang performed data analysis. Zhao Fang and Xieyang Bian performed the optical imaging experiments. Yu Zhang, Zhao Fang, and Xinning Sun performed the MRI experiments. All authors participated in writing the manuscript. Wuwei Ren, Hao Chen and Guoyan Zheng supervised the project.

## Abstract

Preclinical stroke research is severely hampered by the lack of high-throughput, quantitative perfusion imaging tools capable of bridging the gap between functional hemodynamics and structural injury. The emerging second near-infrared (NIR-II) fluorescence imaging offers superior spatial resolution and tissue penetration, but its utility remains restricted by a lack of standardized analysis protocols and inability to resolve in-depth infarct regions. Here, we introduce FIMPA, an integrated analytical framework for standardized, multi-parametric stroke evaluation based on NIR-II fluorescence perfusion imaging. FIMPA systematically processes the image sequence through motion correction and atlas registration to generate an anatomically aligned library of 125 vascular and hemodynamic parameters. Statistical screening identifies 67 stroke-correlated features, enabling objective, data-driven quantification of perfusion deficits. To resolve spatial infarct boundaries, we developed FIMPA-SI (FIMPA Stroke Index), a deep learning-based index utilizing a ViT-UNETR framework. By leveraging self-supervised pre-training with MRI data, FIMPA-SI suppresses dominant vascular signals to translate multi-parametric dynamics into anatomically precise stroke severity maps. Validated in tMCAO mice, FIMPA-SI achieves high spatial concordance with MRI and significantly outperforms conventional laser speckle contrast imaging and single-parameter metrics. FIMPA provides a robust solution for accelerating preclinical therapeutics and advancing optical neuroimaging.

## 1. Introduction

Ischemic stroke, characterized by the critical obstruction of cerebral blood flow, remains a primary cause of global mortality and long-term adult disability [1]. In both clinical and preclinical settings, perfusion imaging serves as an indispensable tool by using intravascular tracers to quantify blood flow through the brain parenchyma [2]. Clinically, computed tomography (CT) and magnetic resonance imaging (MRI) are the gold standards for assessing whole-brain vasculature, which identify the infarct core or potentially salvageable ischemic penumbra by mapping hemodynamic parameters such as cerebral blood flow (CBF), cerebral blood volume (CBV), and mean transit time (MTT) [3–5]. However, the translation of these modalities to preclinical small-animal research is severely limited by high operational costs, restricted accessibility, and insufficient spatiotemporal resolution for murine disease models [6]. Consequently, there is an urgent need for accurate, high-throughput, and cost-effective perfusion imaging solutions to evaluate therapeutic efficacy in the preclinical study pipeline.

Optical imaging has emerged as a compelling alternative for preclinical stroke research due to its inherent cost-effectiveness and high throughput. Laser speckle contrast imaging (LSCI) is widely used for its rapidness and non-invasiveness during perfusion mapping, yet it often suffers from poor spatial resolution and limited quantification due to heavy tissue scattering [6, 7]. While confocal or multi-photon microscopy offer high-resolution vascular detail, they are typically hampered by a restricted field-of-view and the requirement for invasive craniotomy [8, 9]. Traditionally, fluorescence imaging (FI) in the first near-infrared window (NIR-I, 700–900 nm) using Indocyanine Green (ICG) has been utilized to monitor real-time perfusion and blood-brain barrier integrity [10]. However, the utility of NIR-I is constrained by significant skull scattering and limited penetration depth. To overcome these barriers, the second near-infrared window (NIR-II, 1000–1700 nm) has emerged as a superior modality, offering reduced photon scattering and deeper tissue penetration for more precise ischemic localization and perfusion assessment [11–13].

Despite the physical advantages of NIR-II fluorescence perfusion imaging, it remains largely a rapid and qualitative assessment tool. Its wider application in preclinical stroke research is still hindered by two critical limitations. First, quantitative analysis remains elusive due to a lack of standardized data processing pipeline and consensus on significant parameters [14]. Unlike CT and MRI, which utilize extensively validated hemodynamic parameters (e.g., CBF and CBV) to identify the ischemic core, fluorescence perfusion often relies on proprietary software with conflicting definitions for hemodynamic indices [10]. Although time-related parameters like MTT and time-to-peak (TTP) have been demonstrated more robust than intensity-dependent metrics [15], environmental heterogeneity (e.g., camera distance, ambient light) [16] and respiratory motion artifacts [17] continue to introduce significant noise that current ad-hoc protocols fail to correct systematically. Second, NIR-II FI typically suffers from a loss of spatial information in depth. Existing FI systems mainly generate plannar images that lack the depth-resolved detail like CT and MRI, because the fluorescence signal from the deeply seated ischemic core is severely attenuated and entangled with the surface signal. Mesoscopic 3D fluorescence imaging successfully resolves volumetric microangiography at 1-2 mm depth using stereovision-based algorithm, but collapses in diffusive light propagation range beyond several millimeters [18, 19]. Macroscopic fluorescence molecular tomography (FMT) enables tomographic mapping of the fluorescence distribution via physics-driven light propagation model [20, 21]. yet is restricted by a pronounced depth-resolution trade-off, leaving deeper ischemic structures invisible or effectively poorly quantified.

To address these challenges, we developed FIMPA, a standardized NIR-II fluorescence perfusion imaging-based multi-parametric analysis pipeline designed for quantitative and spatially resolved ischemic stroke evaluation. This framework moves the field of FI toward data-driven hemodynamic certainty by integrating automated motion compensation to mitigate respiratory artifacts and atlas-based spatial registration to ensure robust cross-subject comparisons. Next, FIMPA identifies dominant contributors to stroke prediction by extracting a comprehensive library of physiologically interpretable features. Finally, by integrating these multidimensional fluorescence parameters with MRI-derived reference labels, we trained a deep learning model to produce a novel spatial-domain stroke index, FIMPA-SI. This approach effectively bypasses the limitations of traditional thresholding and depth-resolution trade-offs by producing a projection of whole-brain ischemic injury that integrates cumulative damage along the depth axis. Ultimately, FIMPA-SI enables precise, region-specific quantification of stroke burden across the entire brain without the need for additional, high-cost imaging modalities.

## 2. Results

### 2.1 FIMPA workflow

We proposed a quantitative assessment framework for ischemic stroke based on NIR-II FI (Fig. 1). For data acquisition, we developed a customized FI system equipped with 808 nm laser (MFL-808, Artemis Intelligent Imaging, Shanghai, China) for fluorescence excitation and an InGaAs camera (NIRvana-LN, Teledyne, USA) for detection (Fig. 1a). Dynamic FI was performed in ischemic stroke model mice and healthy controls after intravenous injection of CH-4T/FBS solution, a NIR-II contrast agent with excitation peak at 750 nm and emission peak at 1055 nm [22]. For each mouse, the data-acquisition was performed for 203.25 s, during which the fluorescence signal reached a plateau. Details regarding FI experiments and animal preparations are provided in Sections 4.1 and 4.2 of the Methods.

**Figure 1:**
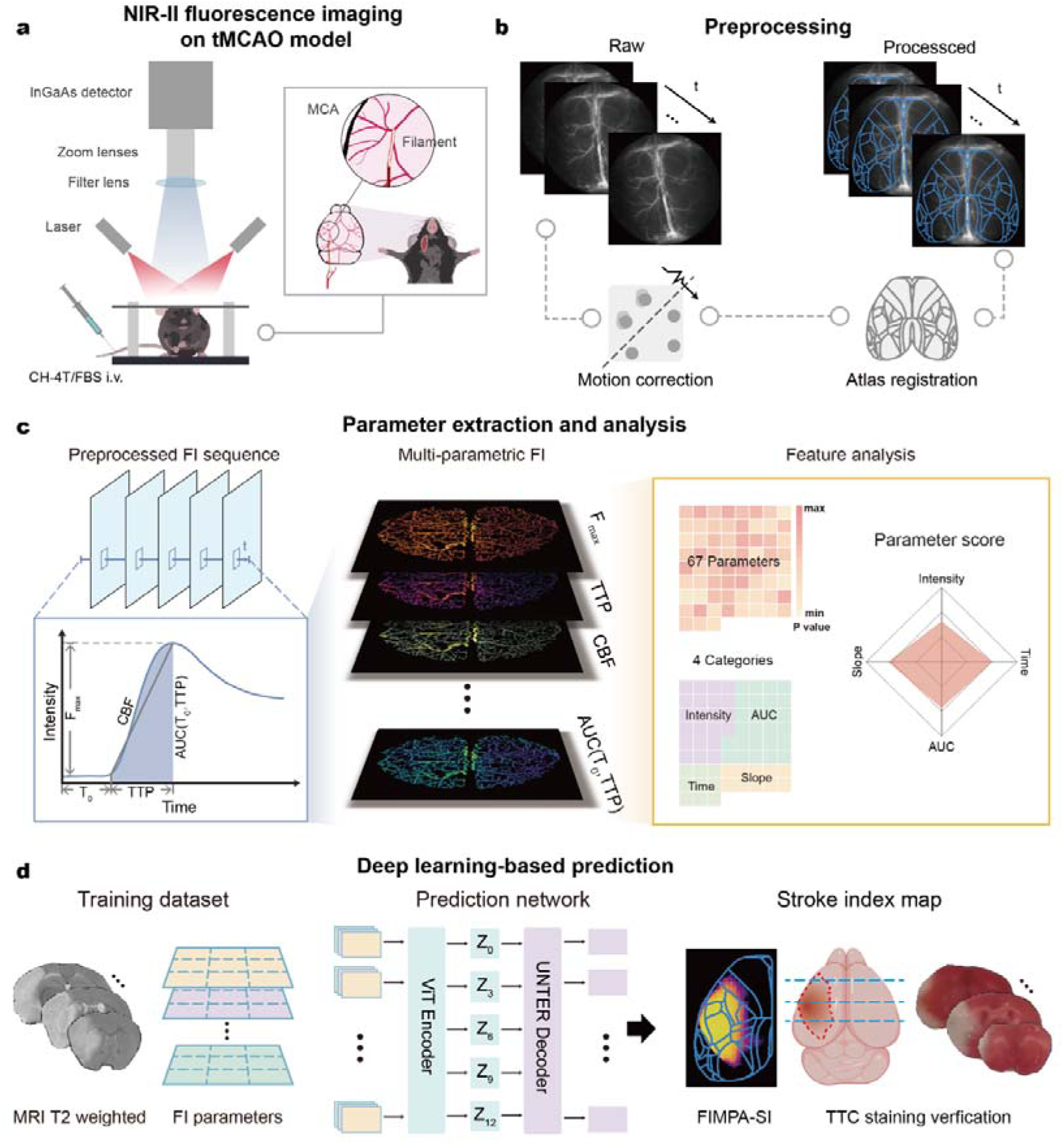
FIMPA: a multi-parametric analysis framework for quantitative stroke evaluation based on NIR-II fluorescence imaging (FI). (a) Dynamic NIR-II FI was performed on tMCAO mice following intravenous injection of the CH-4T/FBS fluorescence probe. (b) The acquired FI image sequence was preprocessed to avoid motion effects and registered with a standard brain atlas prior to parameter extraction. (c) Multiple parametric images (middle) were obtained from the preprocessed image sequence based on the pixel-wise time curve (left). Statistical analysis identifies stroke-correlated parameters throughout the brain (right). (d) FIMPA-SI, a novel stroke index based on a ViT-UNETR framework, indicates the severity of stroke at different brain regions. T2-weighted MRI images were used for training, while TTC-stained slices were used for evaluation.

Following data acquisition, the raw image sequences were preprocessed to remove motion effects using group-wise registration, which ensures frame-to-frame coherence. Then, the registered sequences were mapped into a unified space derived from a standard Brain Atlas [23], enabling cross-subject and multimodal comparisons (Fig. 1b). Next, from the preprocessed sequences, pixel-wise time-intensity curves were computed to derive multi-parametric maps for characterizing vascular architecture and hemodynamic function (Fig. 1c, left/middle). Statistical analysis across the brain identified parameters significantly correlated with stroke (Fig. 1c, right). Finally, an ViT-UNETR framework [24] was designed to generate a new FIMPA Stroke Index (FIMPA-SI), which predicts region-wise ischemic burden based on the extracted parameters in the previous step (Fig. 1d). The learning procedure was trained using atlas-registered MRI T2-weighted imaging (T2WI) images and evaluated against TTC-stained slices, providing spatially resolved quantification of stroke severity directly from FI-derived parameters. This end-to-end pipeline converts raw fluorescence dynamics into quantitative, anatomically standardized assessments. FIMPA-SI relates functional hemodynamics to structural injury, enabling region-specific stroke evaluation without MRI during preclinical research.

### 2.2 Multi-parametric extraction from the FI image sequence

Respiratory motion can introduce imaging artifacts that compromise the accuracy of parameter extraction. Therefore, motion was corrected via group-based registration with a propagated reference frame, using KAZE feature matching to estimate affine transforms for alignment [25, 26] (Fig. 2a). The frame-to-frame coherence has been significantly improved after motion correction, as shown by the pixel-wise intensity-time curve (Fig. 2b), laying a solid foundation for subsequent parametric extraction. All NIR-II FI images were normalized to the 2D Allen Brain Atlas space by landmark-based affine registration using nine manually selected anatomical landmarks, and all parameter extraction and downstream analyses were performed in this normalized space (Fig. 2c). The details of motion correction and atlas registration are referred to Section 4.3 and Supplement. After preprocessing, multi-parametric extraction was performed to quantify vascular morphology and cerebral hemodynamics using various established definitions (Supplementary Table 2). Ischemic stroke has been proven to cause mural cell-mediated cerebral microvessel constriction [27]. Therefore, vascular morphology analysis is an important component of stroke imaging [28, 29]. For morphological analysis, a maximum intensity projection (MIP) image was used as a structural reference to generate the corresponding vascular diameter map (Fig. 2d). To demonstrate the accuracy of the extracted vascular diameter, a square region was enlarged to illustrate segmentation of vessels with varying calibers (Fig. 2e). The cross-sectional profiles of two representative vessels are drawn (V1 and V2), suggesting that even small-sized vessels with a diameter of ∼0.04 mm can be identified and resolved in the diameter map (Fig. 2f). Quantitative validation showed that the full width at half maximum (FWHM) values measured on the MIP were 117.5 μm and 40.1 μm, respectively, closely matching the diameter map (117.6 μm and 39.6 μm). The results demonstrate the accuracy of morphological processing, particularly for small or low-contrast vessels that are challenging to detect in the MIP alone.

**Figure 2:**
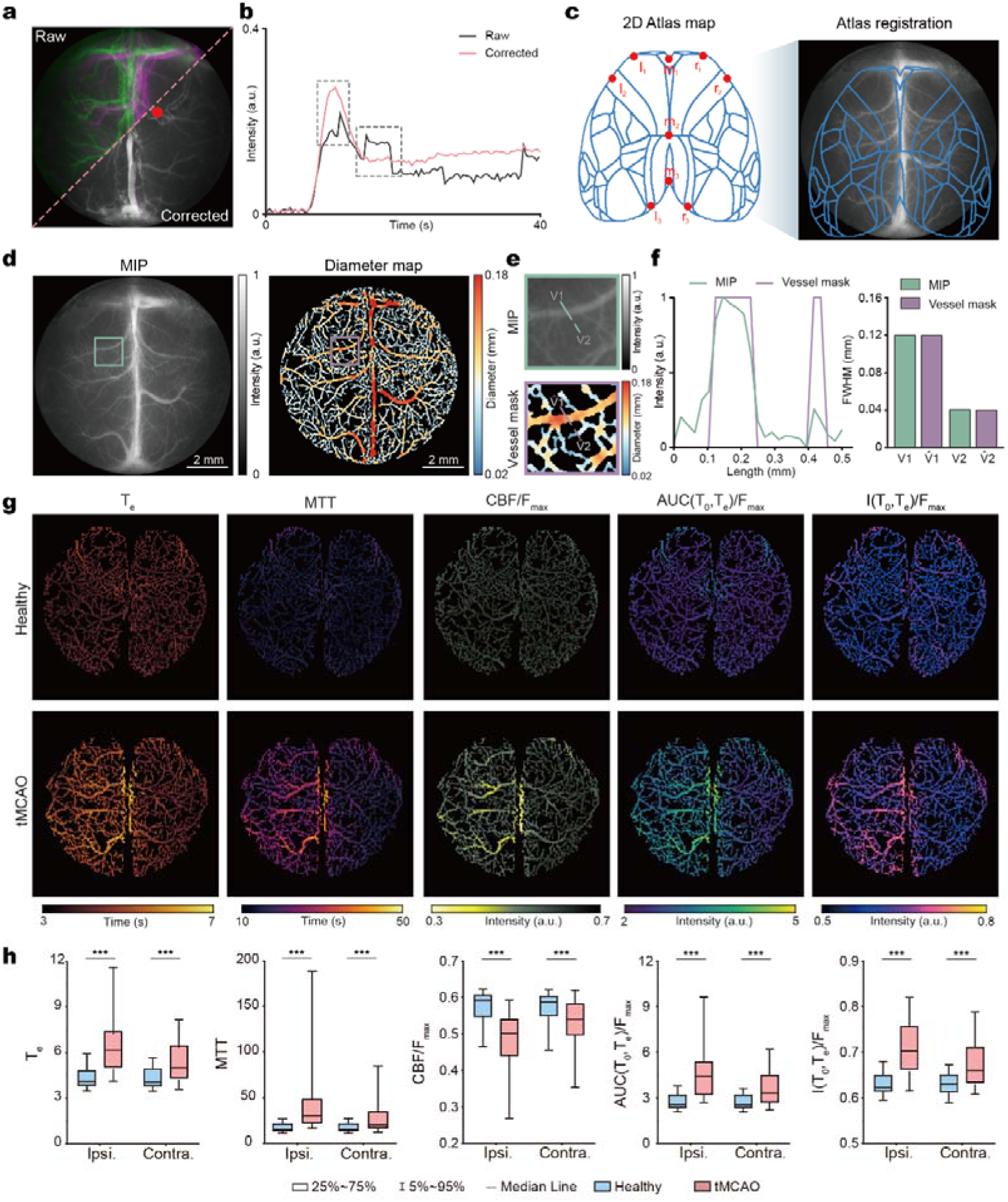
Results of motion correction, atlas registration, and parameter extraction. (a) Comparison of two overlaid frames at t = 9.98 and 10.69 s out of raw and motion corrected image sequences, respectively. (b) Fluorescence intensity-time curves corresponding to the marked red point in (a). (c) Brain atlas overlaid with maximum intensity projection (MIP) image of the FI sequence. (d) MIP image and the corresponding vascular diameter map. (e) Zoomed-in region marked in (d), with two representative vessels highlighted. (f) Line intensity profiles of the two representative vessels are shown in (e), with the full width at half maximum (FWHM) values shown in the bar graph. (g) Selected functional parameter maps of a healthy mouse and a tMCAO model (T_e_, MTT, CBF/F_max_, AUC(T_0_,T_e_)/F_max_, I(T_0_,T_e_)/F_max_), showing a significant difference between the ipsilateral and contralateral hemispheres. (h) Median values of the functional parameters on the ipsilateral and contralateral hemispheres in healthy mice and stroke mice.

Beyond the morphological information, a variety of functional parameters were computed pixel-wise from intensity-time curves to characterize blood flow dynamics and transport behavior. Representative parametric maps including time-related parameters (transit time, T_e_; mean transit time, MTT), flow-related parameters (cerebral blood flow normalized to peak intensity, CBF/F_max_), and intensity-related parameters (area under the curve during the transit interval normalized to peak intensity, AUC(T_0_, T_e_)/F_max_; mean intensity during transit normalized to peak intensity, I(T_0_, T_e_)/F_max_) are shown in Fig. 2g. In healthy mice, these parameters exhibited minimal hemispheric differences, consistent with symmetric perfusion. In contrast, stroke model mice (tMCAO) showed pronounced asymmetries: time-related parameters (T_e_, MTT) were markedly elevated in the ipsilateral hemisphere, indicating prolonged fluorescence probe retention, which may reflect vascular damage or occlusion. The flow-related parameter (CBF/F_max_) was reduced ipsilaterally, suggesting diminished arterial inflow and hypoperfusion. Intensity transition metrics (AUC/F_max_, I/F_max_) were increased ipsilaterally, consistent with prolonged transit and impaired inflow/outflow kinetics. The group-wise summaries (Fig. 2h) further underscore the discriminatory power of these illustrative metrics. Together, the morphology and functional maps derived from FI image sequences provide a more comprehensive quantitative representation of vascular architecture and hemodynamic function than the raw image data, forming a robust basis for standardized analysis in our FIMPA framework.

### 2.3 Quantitative analysis of stroke with the extracted parameters

A library of 125 candidate morphological and functional parameters was generated via multiparameter extraction. Parameters were screened by comparing the stroke and healthy groups (P < 0.001), and 67 highly significant parameters were retained for quantitative and spatial characterization of the infarct region (Fig. 3a). These selected parameters could be classified into four categories: intensity-, time-, AUC- and slope-related metrics. The profiles of the 67 selected parameters are shown in Fig. 3b and reveal systematic increases and decreases in the stroke model, underscoring the need for metrics from all categories to capture distinct aspects of hemodynamic impairment. Feature importance was estimated using Chi-square tests (Fig. 3c and Supplementary Figure S9), with the five most relevant parameters, CBF/F_max_, MTT, AUC(T_0_, TTP)/F_max_, AUC(T_1/2_, T_e_)/F_max_ and AUC(T_0_, T_e_)/F_max_, spanning different categories. Together, these findings indicate that restricting fluorescence quantification to a small subset of parameters may underrepresent the full hemodynamic deficit in the stroke model. To more intuitively illustrate differences between the stroke and healthy groups and to facilitate practical application, the 67 parameters are summed by category and presented as category-level metrics (Fig. 3d and Supplementary Table S3). Compared with healthy controls, tMCAO mice exhibited significant alterations across all parameter categories. Specifically, the total composite score increased from 1.007 in healthy mice to 1.254 in tMCAO mice, with elevated scores across all four categories, intensity (0.258 to 0.275), time (0.160 to 0.220), AUC (0.361 to 0.442), and slope (0.229 to 0.316), reflecting broad functional alterations post-stroke. The detailed parametric analysis method is referred to Section 4.5.

**Figure 3:**
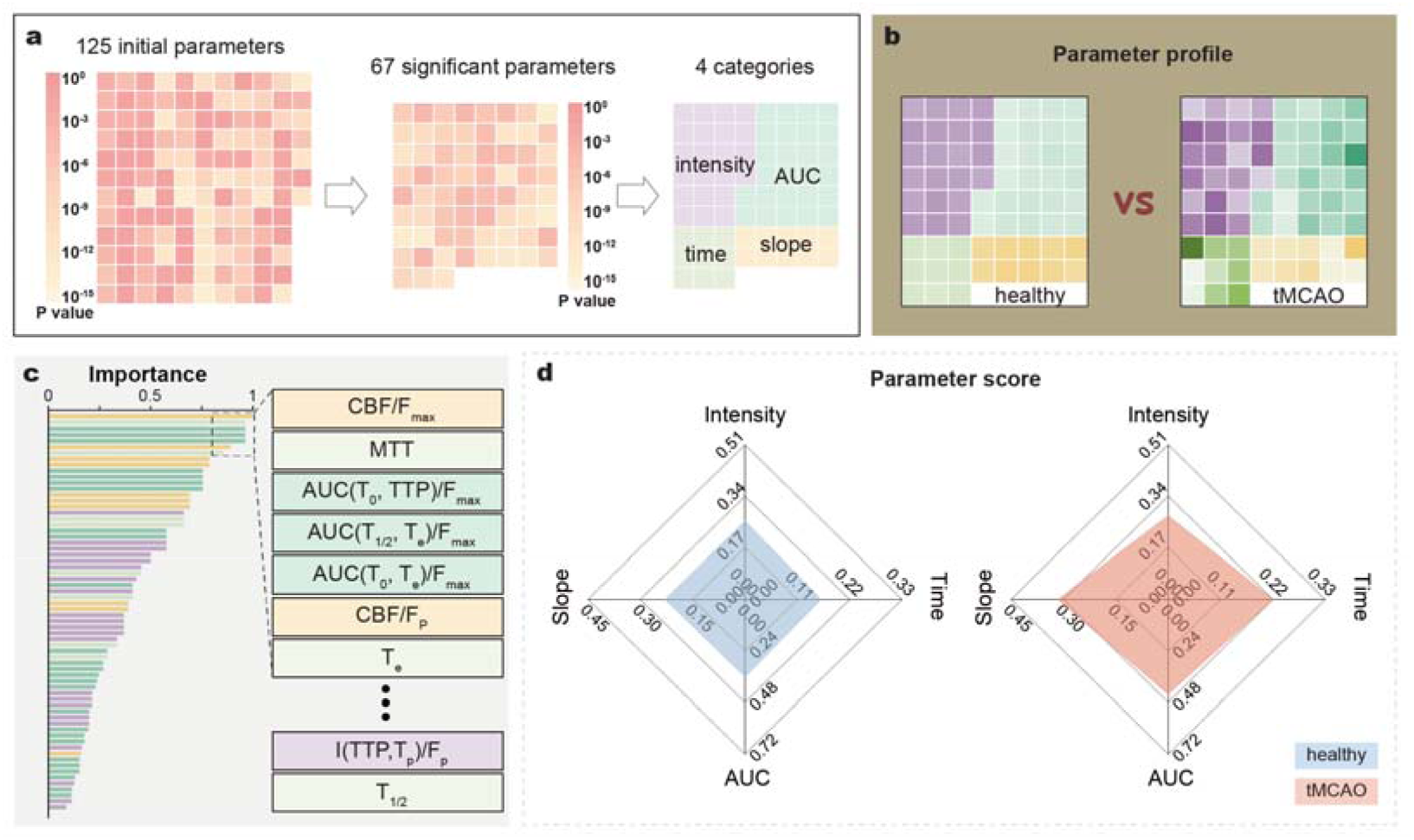
Data significance analysis based on extracted FI parameters. (a) Out of 125 initially extracted parameters, 67 important features were identified with significance tests, and further categorized into 4 classes: intensity-related, AUC-related, time-related, and slope-related. (b) The ratio of mean values between the ipsilateral and contralateral hemispheres for the selected 67 parameters for healthy and tMCAO groups. (c) The rank of parameters according to Chi-square test relative importance. (d) Averaged categorized scores of both healthy and tMCAO mice. The tMCAO group exhibited markedly elevated scores in the time-, slope-, and AUC-related categories compared to healthy controls.

### 2.4 Spatially resolved evaluation of stroke with the stroke prediction network

The extracted FI parameters can effectively differentiate healthy and tMCAO mice. Nevertheless, the precise location and severity of the stroke regions cannot be directly inferred from those parameters. To address this, we have developed a deep learning framework to predict voxel-wise stroke severity maps. The network integrates a Vision Transformer (ViT) encoder and an UNTER decoder (Fig. 4a). The self-supervised pre-training initially enables the network to capture robust contextual representations from the extracted FI parametric features, followed by fine-tuning to map all 67 FI parameters to a transverse plane showing stroke severity distribution with its ground truth (GT) generated from MRI T2WI data. The details for translating the MRI T2WI data to the stroke severity map are given in Section 4.6. This network enables accurate prediction of stroke severity across brain regions solely from FI images. In a representative tMCAO mouse (Fig. 4b), the predicted stroke index (FIMPA-SI) shows strong spatial concordance with the GT, achieving an SSIM of 0.870 and RMSE of 0.066. We thresholded both indices at 25% to delineate stroke boundaries and evaluated the regions of true positives (TP), false positives (FP), and false negatives (FN). A good agreement was found within the lesion core, with minor discrepancies restricted to the periphery (Fig. 4b). A DICE coefficient of 0.92 further confirms precise spatial delineation in this example. Aggregate evaluation across all testing sets (Fig. 4c) shows robust generalization, with a mean SSIM of 0.843 ± 0.022, RMSE of 0.136 ± 0.036, and DICE of 0.833 ± 0.094. The receiver operating characteristic analysis yields an AUC of 0.89, underscoring strong discriminative capability between healthy and stroke regions. To sum up, these results demonstrate that the FIMPA-SI network provides an accurate, spatially resolved assessment of stroke severity at the individual level, maintaining consistent performance across the cohort and effectively bridging quantitative FI metrics with MRI-derived stroke information.

**Figure 4:**
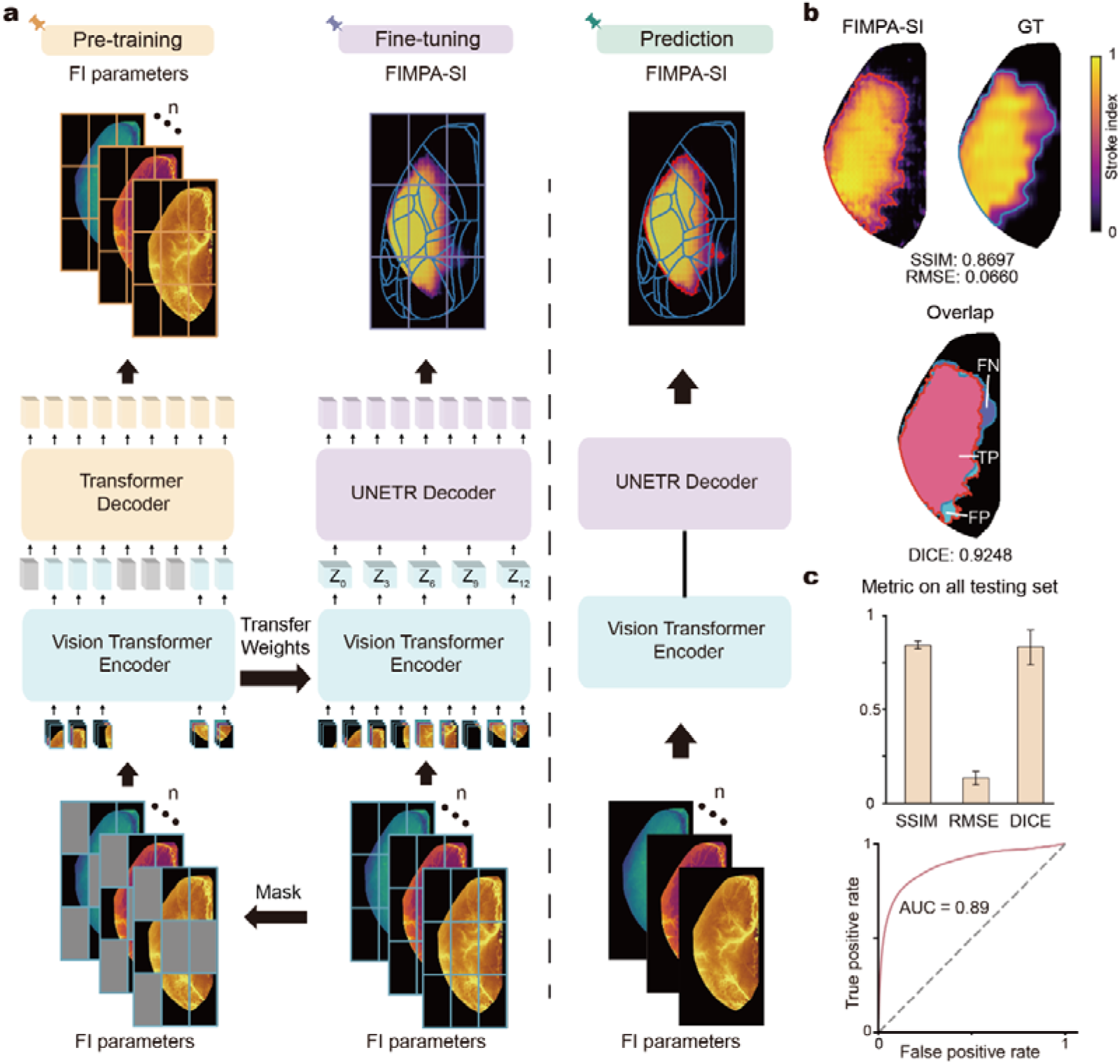
Stroke severity prediction network design and evaluation. (a) The network contains a ViT encoder and a UNTER decoder. Self-supervised pre-training enables the encoder to extract robust latent representations, followed by fine-tuning to map all FI parameters to MRI-based ground truth (GT). Ultimately, the network is capable of predicting stroke severity across different brain regions solely based on FI images. (b) The generated new index (FIMPA-SI) and its corresponding GT were compared by thresholding at 25% to delineate stroke boundaries, and overlaying the regions of true positives (TP), false positives (FP), and false negatives (FN). (c) Quantitative evaluation on all testing sets using different metrics, including SSIM, RMSE, DICE, and ROC.

### 2.5 Comparison of FIMPA-SI to multimodal data

We also compared FIMPA-SI against multimodal data obtained from the same subject. MRI images and TTC-stained slices revealed a pronounced left-hemisphere lesion, with MRI showing marginally larger affected territory than TTC (Fig. 5d). This can be explained by the fact that MRI visualizes both the infarct core and the surrounding affected edematous tissue, while TTC staining specifically marks only the necrotic tissue with loss of enzyme activity [30]. Compared with TTC-stained slices, MRI images were acquired *in vivo* without spatial distortion or misalignment (identical to FI), thus serving as the GT for network training and evaluation.

**Figure 5:**
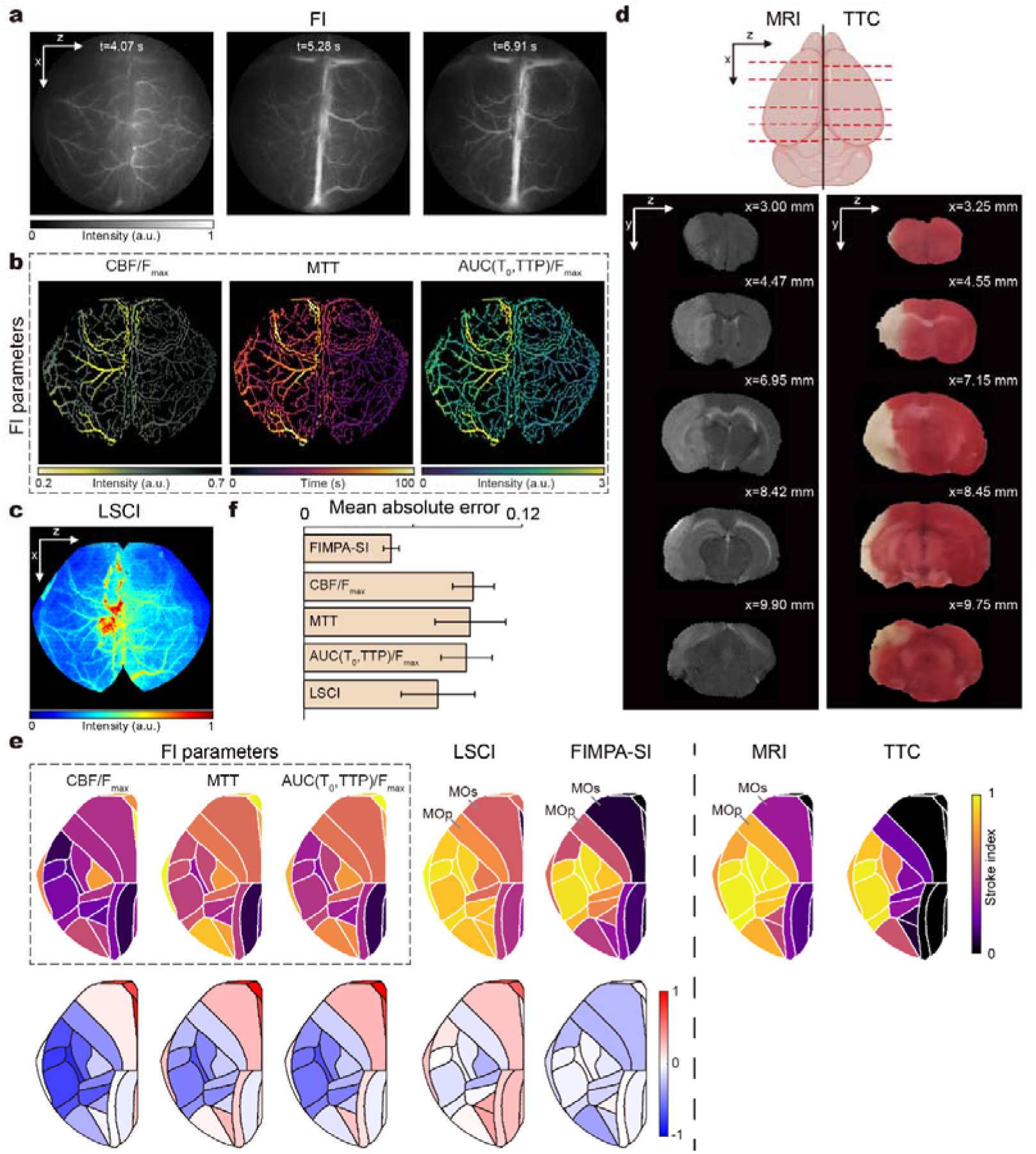
Validation of FIMPA-SI for stroke assessment based on multimodal imaging data. (a) Raw NIR-II FI images recorded at t =4.07, 5.28, and 6.91 s. (b) Top three significant parameters for stroke: CBF/F_max_, MTT, and AUC(T_0_, TTP)/F_max_. (c) A representative LSCI image on the same subject. (d) Paired T2-weighted MRI images and TTC-stained slices. (e) Comparison of the predicted degrees of stroke severity derived from 3 FI parameters, LSCI, and FIMPA-SI. All scores were averaged across segmented atlas regions and compared with stroke indices obtained from MRI and TTC-stained slices. (f) Analysis of relative errors calculated on different methods.

For the illustrative stroke mouse, raw FI images at three time points (t =4.07 s, 5.28 s and 6.91 s) are shown in Fig. 5a. During the initial intensity rise from t = 4.07 s to 5.28 s, the mean value in the affected left hemisphere is significantly lower than that in the right. And this disparity largely disappears once the surface cortical venous signals became pronounced (t = 6.91 s). In Fig. 5b, the three top-ranked FI parameters (CBF/F_max_, MTT, and AUC(T_0_, TTP)/F_max_) sensibly capture these differences: CBF/F_max_ is visibly reduced in the lesion region, while MTT and AUC(T_0_, TTP)/F_max_ are substantially elevated. Similarly, LSCI shows a marked decrease in intensity in the stroke hemisphere (Fig. 5c). While these parameters and modalities enable identification of global signal changes associated with stroke, they are not capable of delineating the precise boundaries of the stroke region.

For quantitative evaluation, all metrics were normalized and averaged over the segmented regions of the Brain Atlas (Fig. 5e, upper). Error maps were computed by calculating the difference between each metric and the MRI GT (Fig. 5e, lower). Across most regions, the errors associated with the single FI parameters are notably larger and more widespread, showing substantial deviation from the true stroke pattern. LSCI, while showing smaller overall errors than the FI parameters, still lacked spatial precision. Critically, it was unable to resolve boundaries that are clearly visible on the MRI, such as the interface between the adjacent regions, the primary motor area (MOp) and the secondary motor area (MOs). In comparison, FIMPA-SI exhibits markedly lower errors, and the difference map closely resembles MRI GT. For tMCAO mice (n = 4) with LSCI, FIMPA-SI achieved the lowest mean absolute error (0.048 ± 0.004), outperforming LSCI (0.093 ± 0.011), CBF/F_max_ (0.092 ± 0.019), MTT (0.089 ± 0.014), and AUC(T_0_, TTP)/F_max_ (0.074 ± 0.020) (Fig. 5f). Collectively, multimodal verification confirms that the proposed index, FIMPA-SI, is capable of linking the functional hemodynamic information with stroke regions by integrating multi-parametric dynamic FI information. FIMPA-SI outperforms any single parameter extracted from the FI image sequences and the conventional LSCI image in spatial accuracy, closely aligning with the MRI-derived stroke pattern.

## 3. Discussion and conclusion

NIR-II FI provides unique advantages for cerebral stroke evaluation in preclinical studies [13, 31, 32], yet its wider adoption has been hampered by two critical challenges: the lack of standardized quantitative analysis protocols [15] and the inability to directly map perfusion features to spatially resolved stroke burden [33, 34]. FIMPA tackles these challenges via an integrated analytical framework that systematically extracts and ranks a comprehensive panel of vascular and hemodynamic parameters, while leveraging deep learning to translate FI-based perfusion features into an anatomically precise stroke index. Our results reveal that this standardized, multi-parametric approach greatly improves the objectivity and spatial accuracy of stroke assessment compared to single FI parameters and LSCI, bridging the gap between FI and stroke-affected structural injury.

A key innovation of our approach is the standardized extraction and evaluation of a broad set of NIR-II FI perfusion parameters for stroke analysis. First, we built a comprehensive parameter library based on preprocessed data that are free of motion effect and unified in a common atlas space. Our pipeline enables reliable, quantitative cross-subject comparisons. Statistical tests confirm that these FI parameters, especially AUC- and slope-related ones, effectively distinguish healthy and stroke subjects, providing clear, objective markers of stroke severity (Fig. 3d). Secondly, we developed a deep learning framework that directly visualizes and quantifies stroke-affected regions. Traditional FI analyses only indicate hemodynamics-related information, where the temporal variation of the fluorescence signals is dominated by the intensity changes from large blood vessels, hiding those subtle changes indicative of tissue infarction in adjacent areas. This presents a significant obstacle to precise spatial assessment of stroke burden. To address this, we designed a self-supervised learning pipeline that combines Masked Autoencoder (MAE) [35] pre-training with a ViT-UNETR architecture [24]. The MAE facilitates robust visual representation learning via masked patch reconstruction, enabling the model to distinguish subtle ischemic tissue signals from the overwhelming fluorescence of large vessels. The ViT-UNETR network combines a vision transformer encoder, capable of capturing long-range spatial relationships and global multiscale features within cerebral vascular networks, with a CNN-based decoder that preserves localized details essential for precise delineation of stroke-affected regions. This synergistic design yields FIMPA-SI that show high spatial concordance with MRI ground truth and TTC staining results, significantly outperforming single FI parameters and conventional LSCI in delineating infarct core and boundaries (Fig. 5e). Collectively, our network enables reliable, region-specific discrimination of ischemic tissue, even in areas with strong vascular signals, enabling interpretable and objective stroke evaluation at high spatial resolution.

Despite the advantages mentioned above, our framework has several limitations. The accuracy and generalizability of FI-based stroke quantification are also influenced by many factors, such as fluorescent probe selection, injection protocol, and imaging systems, all of which may vary across laboratories [16, 36]. It will be interesting to apply and validate our FIMPA framework in diverse experimental conditions. Besides, our imaging experiments focus on the perfusion dynamics of the injected fluorescence probe. The performance of FIMPA on longitudinal studies leveraging the molecular specificity of any targeted fluorescence probes [37] will be highly attractive. Although adaptation of the current protocol would be needed, the basic steps, such as preprocessing, feature extraction, statistical analysis, and learning-based prediction, are expected to remain. Finally, we need to point out that such a standardized image analysis framework based on FI can be expanded to other diseases beyond stroke, such as neurodegenerative diseases [38] and brain tumors [39], where NIR-II FI has been extensively applied.

In summary, FIMPA establishes a robust foundation for the standardized, quantitative evaluation of ischemic stroke using NIR-II fluorescence imaging. By integrating multi-parametric feature extraction with anatomically aligned deep learning prediction, our approach addresses longstanding barriers in fluorescence imaging standardization and objective region-specific stroke analysis. This advance not only bridges the gap between complex hemodynamics and gold standards (MRI and TTC-staining slices) but also enables scalable, high-throughput assessment of ischemic stroke across the preclinical and translational pipeline. FIMPA holds strong potential to accelerate drug development and advance optical imaging-based tools for stroke research and therapeutics.

## 4. Methods

### 4.1 Animal Experiment

All animal care and operation procedures were approved by the Institutional Animal Care and Use Committee (IACUC) at Shanghai Institute of Material Medica, Chinese Academy of Sciences. The study utilized a total of 137 male C57BL/6J mice, 6-8 weeks old and weighing 18-22 g. All animals were housed under a 12-hour light/dark cycle with food and water provided ad libitum. These mice were divided into two groups: 29 animals in a healthy control group and 102 in a disease model group subjected to transient middle cerebral artery occlusion (tMCAO), a widely used model that induces focal cerebral ischemia followed by reperfusion, mimicking key aspects of human ischemic stroke. In this procedure, C57BL/6J mice were anesthetized with 1.5% v/v isoflurane through a mask, and the core body temperature was maintained at 36.5 ℃ using a heating pad. After the neck was shaved and disinfected, a midline neck incision was made, and the soft tissues were bluntly dissected to expose the common, internal, and external carotid arteries. Then, the suture (YuShun Biotechnology Co., Ltd., China) was inserted into the external carotid artery, and carefully introduced into the middle cerebral artery through the internal carotid artery. After a certain duration of occlusion, the suture was withdrawn to achieve reperfusion. Transferred the mice to a clean cage when awake. The model of different infarct volumes was realized by controlling the length of the occlusion time (45-90 min).

### 4.2 *In Vivo* NIR-II Fluorescence Imaging

All *in vivo* fluorescence images were acquired with a homebuilt NIR-II FI system. The illumination module contains an 808 nm laser (MFL-808, Artemis Intelligent Imaging, Shanghai, China). The detection adopts a high-resolution InGaAs camera (NIRvana, Teledyne, USA) with 640 × 512 pixels capable of cooling to -190 ℃ to ensure high signal-to-noise ratio (SNR). The optical path is equipped with a 1150 nm long-pass filter (IR Longpass Filter Kit, Thorlabs, USA) and a lens assembly (Resolv4K, Navitar, USA) for image acquisition and adjustable magnification. For *in vivo* brain imaging, the FOV was set to 12.80 × 10.24 mm^2^. During *in vivo* imaging, the mouse was anesthetized with 1.5% v/v isoflurane through a breathing mask. The scalp was incised to expose the cranium. Sterilized fixation was applied to the mouse skull with tissue glue (1469SB Vetbond, 3M, USA). For both the control and tMCAO groups, each mouse was intravenously injected with 100 μL of a 3 mmol/mL CH-4T/FBS solution via the tail vein during the imaging experiment. The synthesis of CH-4T and the formulation of the FBS solution are described in [22]. Dynamic fluorescence imaging was performed for approximately 3 minutes, with an exposure time of 50 ms, corresponding to a 2.46 fps frame rate and a readout time of 356.502 ms. The temporal image sequence of each perfusion trial contains 500 frames.

### 4.3 Preprocessing of raw image data

Following data acquisition, image registration was performed to correct the motion-induced displacement. A group-based registration strategy was applied to ensure robustness and efficiency. Specifically, 25 consecutive frames were grouped with the first frame assigned as the fixed reference. Afterwards, all subsequent frames within the group were registered to this fixed image. More specifically, all feature points in images were identified using KAZE [25], followed by estimation of the affine transformation for precise alignment [26]. Once intra-group registration was completed in the first group, the last-registered frame was set as the reference for the first image of the next group. This process was iteratively repeated until all frames of the sequence were registered (see Supplement Section 1 for more details).

To facilitate spatial alignment and quantitative analysis, a 2D atlas space was established based on the Allen Mouse Brain Atlas [23]. For high-throughput NIR-II FI datasets, all images were spatially normalized into this atlas space to enable cross-sample comparisons. Image registration was performed by manually identifying nine anatomical landmarks [40] on both the MIP image and the brain atlas. An affine transformation was performed based on the landmarks to align each image to the atlas space. All subsequent parameter extraction and analyses were conducted in the normalized atlas space (see Supplement Section 2 for more details).

### 4.4 Extraction of morphological and functional parameters

Vascular segmentation was performed on MIP images. Tubular structures were first enhanced with the Frangi filter [41], followed by adaptive thresholding to generate a binary vessel mask, denoted as *V*. The corresponding vessel skeleton, denoted as *S*, was subsequently derived from *V* [42]. The local vessel radius *r*(*x*, *y*), can be computed as follows:

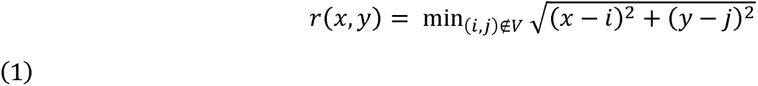

For the arbitrary position (*x, y*) ∊ *S*, *r*(*x*, *y*) represents the minimum Euclidean distance from the local point to the boundary of *V* [43]. The diameter map was obtained by averaging overlapping provisional assignments for each pixel (Supplement Fig. S3a). The detailed implementation of the diameter calculation is given in Supplement Algorithm S1.

Before parameter extraction, each image was denoised with a Gaussian filter (σ = 1) and then filtered with a Savitzky-Golay filter [44]. Various hemodynamic parameters were derived from these intensity-time curves (Supplement Figure S3b), which were further categorized into four groups: 1) time-related parameters such as time to peak intensity and probe clearance time; 2) intensity-related parameters comprising values at critical time points; 3) slope-related parameters reflecting fluorescence change rates; and 4) area under the curve (AUC)-related parameters calculated by integrating an intensity-related parameter over a specific interval (Supplement Table 2).

### 4.5 Statistical analysis of the extracted parameters

To evaluate the relationship between the extracted functional parameters and stroke outcomes, we performed a comprehensive statistical analysis. Normality was assessed using the Shapiro–Wilk test [45], and homogeneity of variance was evaluated using the F-test. For between-group comparisons, Student’s t-test was used when assumptions of normality and equal variances were met; Welch’s t-test was used when variances were unequal; and the Mann–Whitney U test was used when normality was not satisfied. Statistical significance was defined as P < 0.05. Subsequently, all extracted parameters were ranked according to their importance based on the Chi-square test results. All parameters were normalized by calculating the ipsilateral-to-contralateral hemisphere mean ratio. Since certain parameters, such as slope-related parameters, typically show physiologically lower values in the ipsilateral hemisphere post-stroke, we took the reciprocals of their normalized ratios to ensure that higher values consistently reflected abnormality. For example, if a parameter decreased post-stroke (resulting in a ratio < 1), its reciprocal (>1) ensured a higher value aligned with greater injury severity. Finally, individual parameter scores were calculated by multiplying each importance weight by the corresponding adjusted value. A category total score was then derived by summing relevant individual parameter scores within the category. All analyses were performed in Matlab (2021b, MathWorks, MA, USA) and OriginPro 2024 (OriginLab, Northampton, MA, USA).

### 4.6 Learning-based stroke prediction

Training and validation were performed using multimodal data: MRI T2WI images for training, LSCI for comparative validation, and ex vivo TTC-staining slices for infarction verification. The MRI T2WI images were acquired on a 9.4 T high-field MRI scanner (BioSpec 94/20, Bruker, Germany) using T2-TurboRARE sequence with the following settings: repetition time (TR) = 3138 ms; echo time (TE) = 33 ms; number of slices = 30; slice thickness = 0.5 mm; FOV = 20 × 20 mm^2^; reconstruction matrix = 256 × 256. The LSCI images were acquired on RFLSI III (RWD Life Science, China) with the following settings: acquisition time = 5 s; exposure time = 10 ms; FOV = 11 × 11 mm². For TTC validation, brains were sectioned coronally after extraction, stained with 1% TTC at 37°C, fixed in 4% paraformaldehyde, and infarct volumes were quantified. The ground truth data are generated from MRI T2WI images. First, hyperintense lesion areas were segmented using maximum entropy thresholding [46], with manual correction to exclude non-pathological signals. Each MRI slice was then registered to the 3D atlas space using QuickNII [47]. Next, the segmented stroke masks were mapped to the 3D atlas space and projected along the y-axis to obtain a normalized local volumetric distribution of infarct involvement, named the stroke index (SI) map. This distribution was further interpolated along the x-axis and matched to the corresponding FI images for direct spatial comparison (Supplement Section 4 and 5).

The prediction network employed a dual-stage pipeline comprising self-supervised pre-training and subsequent downstream fine-tuning. During pre-training, we adopted a Masked Autoencoder (MAE) framework [35] with a ViT backbone. This model receives randomly masked FI parameter patches, and the encoder processes only the visible subset, while a lightweight decoder reconstructs the complete input. By compelling the model to reconstruct missing information from limited observations, this approach encourages the encoder to aggregate long-range dependencies and capture the global anatomical context, which is critical for medical image analysis. After pre-training, the decoder is discarded, and the pre-trained ViT encoder is integrated into a ViT-UNETR architecture [24] tailored for stroke severity prediction. In this stage, the transformer encoder captures global and multiscale spatial information from the FI parameters, while a CNN-based decoder, connected via multi-resolution skip connections, facilitates the integration of high-level semantic features and spatially precise information. This configuration enables effective translation of FI parameters into 2D atlas-aligned stroke index maps, supporting detailed, spatially accurate prediction of infarct regions. Implement details are provided in Supplement Section 6.

## Acknowledgment

This work was supported by the Science and Technology Commission of Shanghai Municipality (No. 25JS2830300, Wuwei Ren/Hao Chen; YDZX20233100004032, Hao Chen), the National Natural Science Foundation of China (No. 62105205, Wuwei Ren; No. 82572286, Hao Chen; No. 62471293, Guoyan Zheng), the Strategic Priority Research Program of the Chinese Academy of Sciences (No. XDB1060000, Hao Chen), Shanghai Municipal Science and Technology Major Project, the National Key Research and Development Program of China (No. 2023YFA1800804, Hao Chen), Science and Technology Innovation Key R&D Program of Chongqing (No. CSTB2023TIAD-STX0006, Hao Chen), STI2030-Major Projects (No. 2021ZD0203900, Hao Chen).

## Data availability

Data underlying the results presented in this paper are not publicly available at this time but may be obtained from the authors upon reasonable request.

